# Phase Separation and Correlated Motions in Motorized Genome

**DOI:** 10.1101/2022.05.10.491350

**Authors:** Zhongling Jiang, Yifeng Qi, Kartik Kamat, Bin Zhang

**Affiliations:** Department of Chemistry, Massachusetts Institute of Technology, Cambridge, MA, USA

## Abstract

The human genome is arranged in the cell nucleus non-randomly, and phase separation has been proposed as an important driving force for genome organization. However, the cell nucleus is an active system, and the contribution of non-equilibrium activities to phase separation and genome structure and dynamics remains to be explored. We simulated the genome using an energy function parameterized with chromosome conformation capture (Hi-C) data with the presence of active, nondirectional forces that break the detailed balance. We found that active forces that may arise from transcription and chromatin remodeling can dramatically impact the spatial localization of heterochromatin. When applied to euchromatin, active forces can drive heterochromatin to the nuclear envelope and compete with passive interactions among heterochromatin that tend to pull them in opposite directions. Furthermore, active forces induce long-range spatial correlations among genomic loci beyond single chromosome territories. We further showed that the impact of active forces could be understood from the effective temperature defined as the fluctuation-dissipation ratio. Our study suggests that non-equilibrium activities can significantly impact genome structure and dynamics, producing unexpected collective phenomena.

## Introduction

The eukaryotic genome is organized non-randomly in the three-dimensional space, and genomic regions with varying transcriptional activities occupy well-defined regions.^1–9^ Transcriptionally active chromosomes are frequently found near the nuclear interior,^10^ while heterochromatin tends to reside at the periphery close to the nuclear lamina.^11^ The lack of mixing among different chromatin types is evident from chromosome conformation capture (Hi-C)^12,13^ and single-cell imaging experiments.^14,15^ The contact maps produced from these experiments support that euchromatic regions share more interactions among themselves than with heterochromatin, giving rise to the so-called compartmentalization and the emergence of *A*/*B* compartments. Compartmentalization could arise from the microphase separation of polymeric systems, which has been proposed as a key mechanism for genome organization.^16–26^ Molecular mechanisms driving chromosome phase separation are beginning to emerge as well.

Euchromatin and heterochromatin are known to exhibit distinct chemical modifications^27–30^ and interact with different regulatory proteins^28,29^ and nuclear bodies.^9,31–33^ The chemical modifications themselves could drive chromatin demixing by altering nucleosome interactions. For example, acetylation could neutralize the positive charges on histone tails to weaken their binding with DNA, ^34–37^ while methylation may lead to under-twisted DNA that strengthens its interactions with histones. ^38,39^ It’s conceivable that nucleosomes with similar histone marks share more favorable interactions, driving their separation from other differentially modified ones. On the other hand, chromatin regulators associated with specific histone marks often contain intrinsically disordered regions that exhibit multivalent interactions to undergo liquid-liquid phase separation themselves. ^40–46^ These proteins may further contribute to the compartmentalization of various chromatin types.

Besides passive interactions, phase separation can also arise due to the many nonequilibrium processes inside the cell nucleus. ^47–49^ These processes, including transcription, chromatin remodeling, etc., ^50,51^ often consume ATP to drive conformational changes in molecular motors and exert forces on chromatin. Active forces break the detailed balance to produce out of equilibrium genome organizations beyond those dictated by passive interactions. Pioneering studies from the Menon group have shown that active processes could lead to non-random chromosome positions.^47,52^ However, the impact of non-equilibrium activities on the compartmentalization of different chromatin types has not been investigated. Since they are concentrated in euchromatic regions, active processes may produce a temperature gradient between euchromatin and heterochromatin,^53^ contributing to their phase separation.^48,54^ Active and passive mechanisms may both influence genome structure and dynamics, but their combined effect remains to be explored.

We carried out Brownian dynamics simulations to evaluate the role of passive interactions and active molecular motors on genome organization. A potential energy function derived from Hi-C data was used to account for effective interactions that fold individual chromosomes and drive their phase separation. Non-equilibrium activities were added to euchromatic regions by introducing nondirectional active forces. Computer simulations revealed a striking impact of active forces on genomic phase separation. While both passive interactions and active forces facilitate chromosome phase separation, they position *B* compartments in opposite directions. Competition between the two mechanisms produces genome organizations seen in normal and inverted nuclei. In addition to genome structures, the simulation framework naturally allowed the quantification of chromosome conformational dynamics. We observed that active forces with random orientations give rise to spatial correlations between genomic segments that are several micrometers apart on length scales comparable to those observed experimentally. Our study suggests that non-equilibrium activities could give rise to novel collective behaviors and provide new insights into genome organization.

## Results

### Computational model for the motorized genome

We introduced a coarse-grained model for the diploid human genome to approximate the role of loop extrusion^55,56^ and phase separation^16–19,21,22^ on genome organization effectively. We modeled the genome at the one megabase (MB) resolution as a collection of 46 chromosomes inside spherical confinement (Figure 1). Each chromosome is represented as a string of one-megabase-size beads, which can be assigned as one of three types, *A, B*, or *C*. *A* and *B* correspond to the two-compartment types that contribute to the checkerboard patterns typically seen in Hi-C contact maps,^12,57^ and *C* marks centromeric regions. The compartment assignments provide chemical specificity for the coarse-grained beads to model specific interactions and capture microphase separation. In addition to the block co-polymer setup, we introduced intra-chromosome interactions that vary as a function of the sequence separation between two genomic segments. This “ideal” potential approximates the crosslinks produced by Cohesin molecules via their extrusion along chromatin, promoting chromosome territory formation. ^24,58,59^ Detailed expressions of the energy function can be found in the Supporting Information. Interaction parameters in the ideal and compartment potential were tuned to reproduce various average contact probabilities determined from Hi-C experiments for GM12878 cells using the maximum entropy optimization algorithm.^58,60–62^

**Figure 1:**
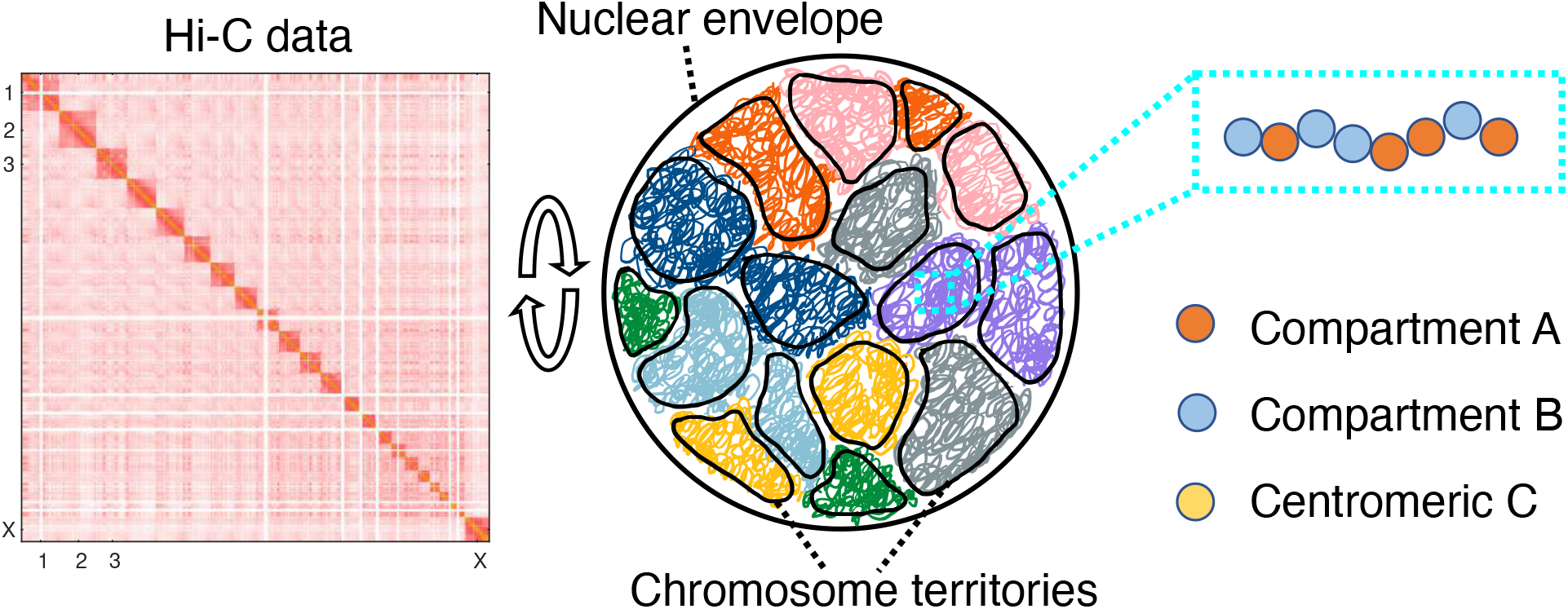
Illustration of the genome model parameterized with Hi-C data. The human genome consisting of 46 chromosomes was modeled in a spherical confinement of 10 *μ*m in diameter. Each chromosome was represented as a string of coarse-grained particles that are one MB in size. We used Hi-C data to identify each bead as compartment *A* (orange), compartment *B* (blue), or centromeric region *C* (yellow) and to parameterize the interaction energy among them.

We carried out Brownian dynamics simulations of the coarse-grained model to produce an ensemble of genome structures (see *Methods Section* for simulation details). These structures support the formation of territories, and individual chromosomes exhibit collapsed conformations with minimal intermingling among them (Figure S1). We further determined an in-silico contact map by averaging over 3D structures to compare with Hi-C data. Given the genome model’s coarse resolution, the simulated contact map inevitably misses fine structural features such as chromatin loops^57^ and topologically associating domains.^63,64^ Nevertheless, it exhibits the typical plaid patterns that arise from the phase separation of *A*/*B* compartments (Figure S1).

An important feature of genome organization that’s amiss in the simulated structures is the peripheral localization of *B* compartments. As shown in the radial density profiles (Figure 2a), a significant amount of *B* compartments resides near the nuclear center. We note that since parameters were uniquely derived from Hi-C data, the observed mischaracterization of genome organization is due to the model’s inherent design, i.e., the specific functional form of the energy defined in Eq. S1, rather than a lack of parameter fine tuning. Indeed, as recognized in previous studies, explicit interactions with nuclear lamina, ^19,32,65,66^ separate treatment of intra- and inter-chromosome interactions,^17^ or assuming repulsive interactions between the compartments, ^18^ might be necessary to position *B* compartments towards the nuclear envelope.

**Figure 2:**
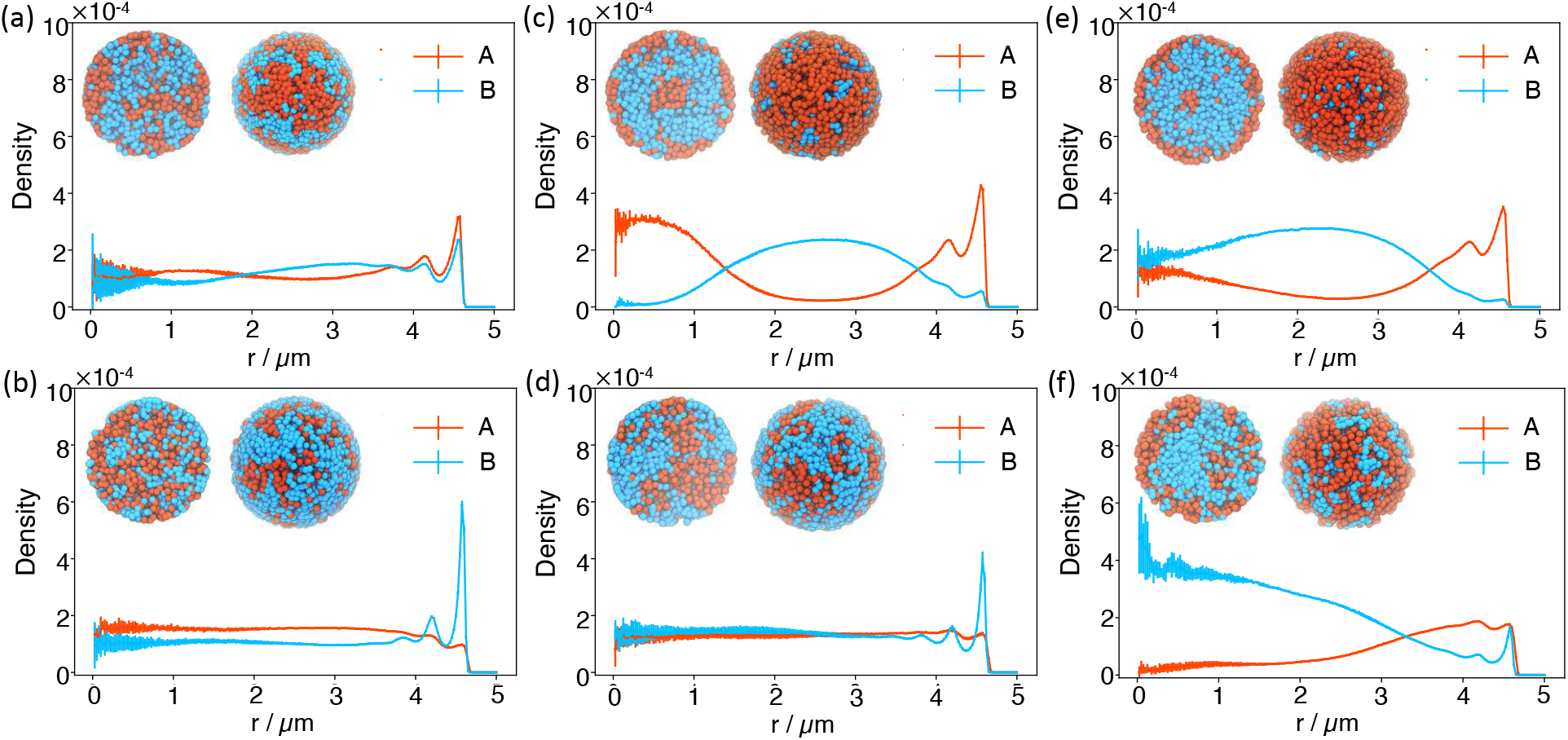
Radial distribution profiles of *A/B* compartments at different interaction strengths between *B* compartments with (top) and without (bottom) the presence of active forces. The interaction strength between *B* compartments was increased by two and three folds from left to right. The insets provide two different views of a representative configuration for each system. The left image corresponds to the front view of a cross-section of the genome sliced through the nuclear center, while the right image corresponds to the view from the nuclear envelope. In these images, *A/B* compartments are colored orange and blue, respectively.

While the model does not fully capture all features of genome organization, it stands out due to the simplicity of its design. The model serves as a valuable baseline to explore mechanisms in addition to loop extrusion and compartmentalization for genome organization. ^67^

### Active forces drive peripheral localization of heterochromatin

The cell nucleus is an active system enriched with ATP-driven processes. The ideal potential was specifically designed to account for the role of molecular motors like Cohesin that can compact individual chromosomes.^55,56^ Other active processes, including transcription and nucleosome remodeling,^51,53,68–71^ can also lead to non-conservative forces to perturb chromosome structures. The effect of these processes, together with passive interactions arising from epigenetic modifications and chromatin regulators, may produce the effective compartment specific potential inferred from Hi-C data.

To disentangle the role of passive and active forces on genome organization, following prior studies,^47,72–74^ we model the effect of transcription as self-propelled particles with random, nondirectional noises. For example, in addition to the typical random forces in the Brownian motion that satisfies the fluctuation-dissipation theorem, an extra term, ***f**_i_*(*t*) was added to the equation of motion for particle *i* at time *t*. The active forces have zero mean, 〈***f**_i_*(*t*)〉 = 0 and the correlation for forces at different times is

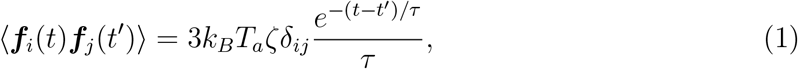

where *ζ* is the friction coefficient and *k_B_* is the Boltzmann constant. *k_B_T_a_* corresponds to the energy released by force generating events and *τ* accounts for the finite lifetime of these events. Active forces were only applied to *A* compartments, i.e., euchromatin, which consist of approximately 40% of the genome.

We first studied genome structures with the presence of non-correlated active forces, i.e., white noises, in the limit of *τ* = 0. In this case, the correlation function becomes 〈***f**_i_*(*t*)***f**_j_*(*t*′)〉 = 6*k_B_T_a_ζδ_ij_δ*(*t* – *t*′). We start with *T_a_* = 10*T*, which is comparable to the amount of energy released from hydrolyzing a single ATP^75^ with *T* as the room temperature. Introducing active forces caused a dramatic change in the radial density profiles. There is a sharp peak for *B* compartments now at the nuclear periphery (Figure 2b), resembling the formation of lamina-associated domains (LADs).^76^ Accompanying the shift of *B* compartments is the increase of *A* compartments near the center, further driving the interior localization of chromosomes with more *A* compartments (Figure S2). As shown in Figure S3, these results are robust with respect to the strength of active forces. Restricting the activities to only the top half of most active *A* compartments also produced similar trends (Figure S4).

### Balancing passive and active forces for heterochromatin localization

In our model, *B* compartments experience attractive interactions among themselves. Attractions could arise from direct contacts between histone proteins and the DNA^44,77,78^ or mediated by chromatin regulators, including HP1, PRC1, and PRC2,^40,41,79–81^ that promote multivalent interactions. To explore the combined role of passive and active forces on genome organization, we carried out additional simulations at different interaction strengths between *B* compartments with varying magnitudes of active forces.

As shown in Figures 2c and 2e, enhancing the interaction between *B* compartments by a factor of two or three causes the opposite effect on genome organization compared to adding active forces. The density profiles are in stark contrast to the one produced by the original interaction strength. Heterochromatin in the new simulations is closer to the nuclear interior, resulting in genome organizations that resemble the inverted nuclei. Strong interactions among heterochromatin have indeed been proposed to drive the formation of inverted nucleus observed in neuronal cells. ^82^ Again, introducing active forces aligns *B* compartments towards the nuclear envelope (Figures 2d and 2f), though the effect becomes less dramatic at very strong passive interactions.

The competition between the two forces in positioning heterochromatin is summarized in a phase diagram (Figure 3a). We defined a collective variable that quantifies the radial distribution of *B* compartments using the ratio of the two average densities computed for regions close to and far away from the nuclear envelope. We separated the two regions with a distance cutoff of 3.85 *μm*. The variable increases for larger active forces and weaker passive interactions, supporting their counter effects on genome organization. We further classified systems with variable values larger than 0.65 and less than 0.2 as normal (blue) and inverted (red) nuclei, respectively based on the visual inspection of the simulated structures.

**Figure 3:**
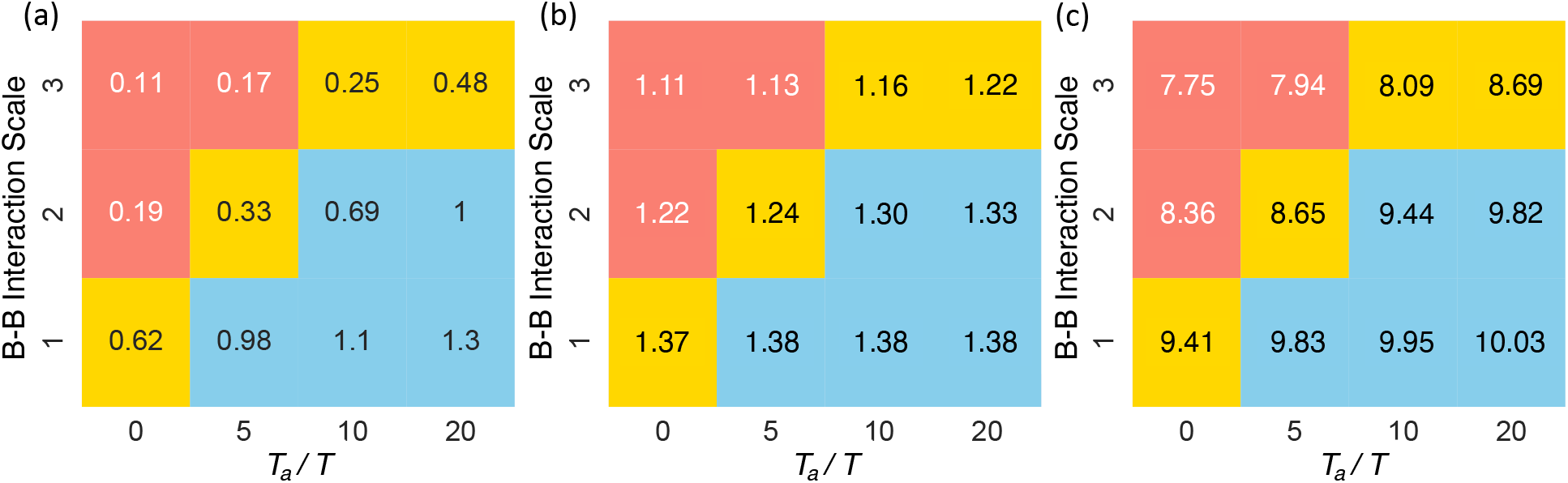
Phase diagram of genome organization. Three phases correspond to the conventional (blue), intermediate (yellow), and inverted (red) nuclei. The *x* and *y* axes indicate the strength of active and passive forces. The number in each bin measures the average density ratio of the outer shell over the inner sphere (a), the average volume assigned to each heterochromatin region (b), and the radius of gyration for the largest cluster formed by *B* compartments (c). See also Figure S7 for the corresponding results for *A* compartments.

To provide a more detailed characterization of the impact of active and passive forces on 3D genome organization, we performed Voronoi tessellation of the simulated structures (Figure S5). Through the tessellation, we identified genomic regions that are nearest neighbors in space and connected them to determine the largest connected component network for *B* compartments. We found that the networks include almost all *B* compartments in all setups (Figure S6), supporting the role of both active and passive forces in promoting phase separation. The average volume assigned to each *B* compartment is also relatively conserved with respect to parameter changes (Figure 3b). However, the radius of gyration of the network showed dramatic variations, indicating changes in the morphology and surface area of the phases (Figure 3c). Increasing the interactions among *B* compartments compacts the networks, while active forces applied to *A* compartments grow their branches.

### Active forces enhance correlated motion among chromatin

In addition to affecting steady-state distributions, non-equilibrium activities are known to give rise to emergent dynamical behaviors absent in equilibrium systems. ^83^ Chromosomes have indeed been reported to exhibit coherent motions across length scales beyond individual territories as a result of ATP-driven processes. ^84–87^ Whether the so-called scalar noise, i.e., the ones studied here with no orientational preference, contributes to such coherent motion has been controversial. Several studies have suggested that scalar noises cannot significantly impact the dynamical correlation length. ^72,84^ Instead, vector events that produce force dipoles have been proposed as essential for enhancing correlation, possibly via a nematic ordering of chromatin. ^72,84,88^ However, these studies did not explicitly account for specific interactions and chromosome compartmentalization, which have been shown to produce dynamical correlations within individual chromosomes. ^74,89^ We further studied how passive interactions and scalar forces impact whole-genome conformational dynamics.

In agreement with previous studies,^74,89^ we observed dynamical correlations among chromatin segments without the presence of active forces. Following Ref. 74, we characterized genome-wide correlated motions using the displacement correlation function defined as

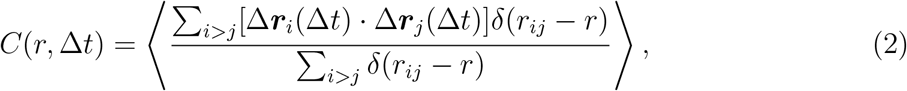

where the angular brackets indicate ensemble averaging and *i, j* index over chromatin segments. Δ***r**_i_*(Δ*t*) corresponds to the displacement of particle *i* over the time interval Δ*t*, · represents the dot-product of the two displacement vectors, and *r_ij_* is the distance between particle *i* and *j*. As shown in Figure 4a, while the correlation function decays quickly as a function of the separation length *r* for small time intervals, it increases significantly for longer Δ*t*. We further defined the correlation length, *l_c_*, using an exponential fit as *e*^−*r/l_c_*(Δ*t*)^ to *C*(*r*, Δ*t*). The dependence of *l_c_* on Δ*t* is shown in Figure 4c.

**Figure 4:**
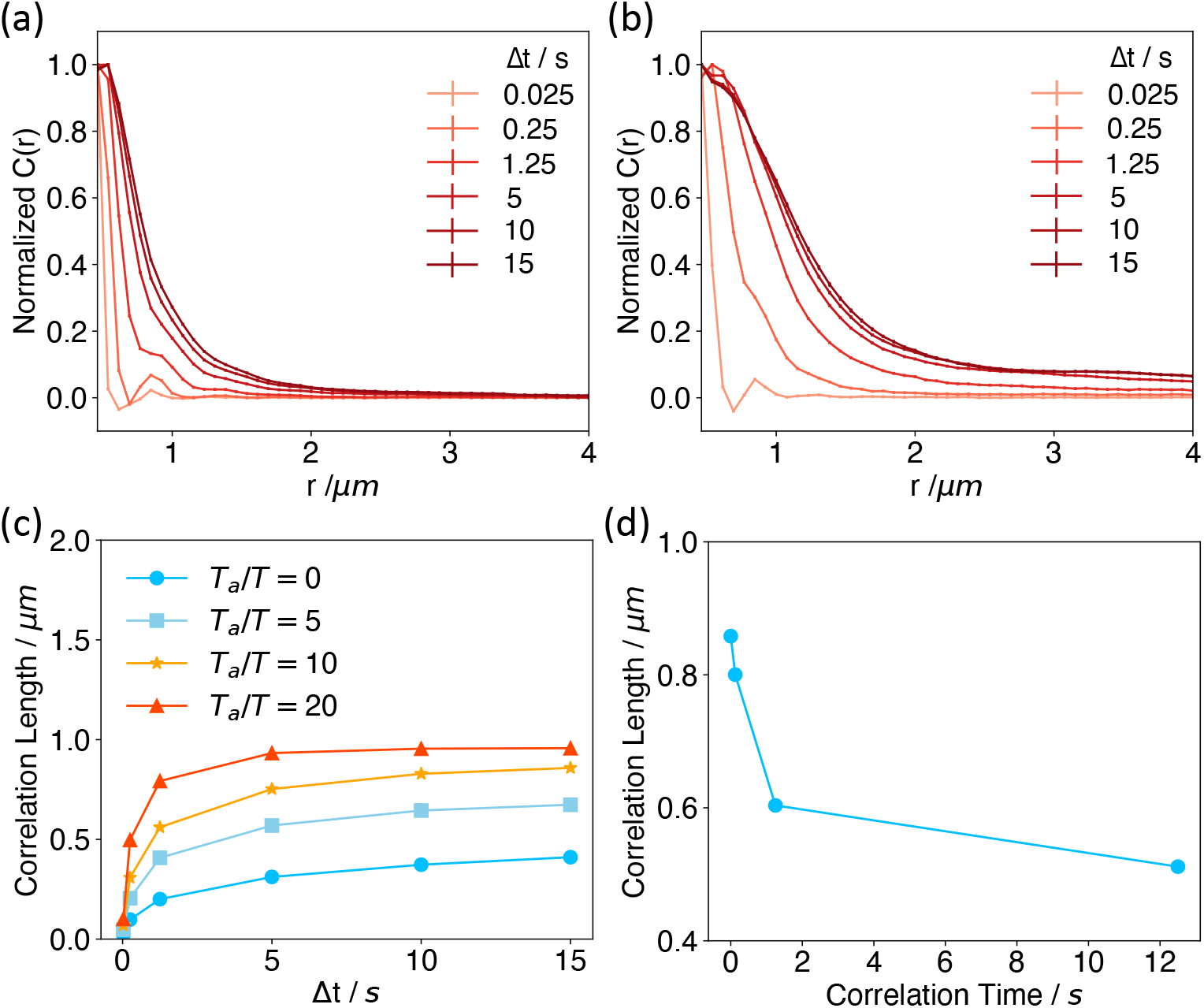
Active forces enhance long-range correlations among genomic loci. (a,b) Displacement correlation functions at various time separations computed without (a) and with (b) the presence of active forces. The correlation functions at different time intervals were normalized such that the maximum values are unity. See Figure S8 for results at different strengths of active forces. (c) Correlation lengths as a function of the time separation for different active forces. (d) Dependence of the spatial correlation length determined at *T_a_* = 10*T* and Δ*t* = 15s as a function of the correlation time of the active forces.

We next examined spatial chromosome correlation in the presence of active scalar forces with *T_a_* = 10*T*. As shown in Figure 4b, a significant increase in the correlation length can be observed, especially for larger Δ*t*. The precise dependence of *l_c_* on Δ*t* at several values of *T_a_* is provided in Figure 4c. These results support a significant role of molecular motors that produce nondirectional forces on chromosome dynamics.

To further break down the intra- and inter-chromosomal contributions, we limited the genomic pairs to those from the same or different chromosomes when computing *C*(*r*, Δ*t*). As shown in Figure S9, active forces enhance correlations both within and across chromosomes. As expected, the correlation length within individual chromosomes cannot go beyond a certain value characteristic of the territory size. However, the inter-chromosomal correlations increase to much larger values.

The results presented so far are for active forces with zero correlation time, i.e., *τ* = 0 as defined in Eq. 1. However, forces at different time intervals could be correlated due to the finite lifetime of chemical processes, producing nonzero values for *τ*. Since precise values for *τ* are not available, we consider a range of values from 0.125 to 12.5 s that roughly matches the rate for chromatin remodeling.^90^ As shown in Figure 4d, the correlation length for Δ*t* = 15s decreases significantly as *τ* increases. In addition, we found that nonzero values for *τ* also reduced the impact of active forces on the peripheral localization of *B* compartments (Figure S10).

### Effective temperature model for motors

We further resorted to the theory of effective temperature ^91^ to provide intuitions for the role of active forces on genome organization. Effective temperature has proven useful for studying non-equilibrium systems and glasses.^92–94^ We quantify the effective temperature with the fluctuation-dissipation ratio^53,92–95^ defined as

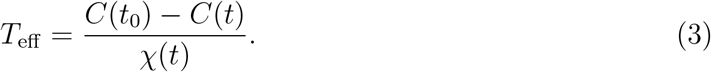

*C*(*t*) represents the correlation function for the single particle density fluctuation with observables defined as 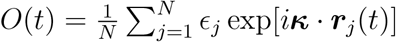 and 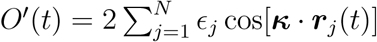, and *χ*(*t*) is the corresponding response function. More detailed expressions for computing the effective temperature are provided in the Supporting Information.

We first analyzed systems with non-correlated active forces at *τ* = 0. In such cases, the equation of motion for *A* compartments is identical to the one without active forces but at an elevated thermal temperature of *T_a_* + *T*. We found that the susceptibility-correlation plots exhibit a single linear regime across all timescales (Figure 5a). The effective temperatures defined from the negative slope of such plots indeed agree with *T_a_* + *T* (Figure 5b). Therefore, our analyses suggest that the overall system is equivalent to *A*/*B* compartments at equilibrium with two separate thermostats of different temperatures. ^47,53,96^ Phase separation of liquid mixtures with differential temperature has indeed been well-documented,^48,73,97,98^ which may explain the changes seen in Figure 2. The higher temperature of *A* compartments makes them entropically favorable to stay in the interior with more conformational space.

**Figure 5:**
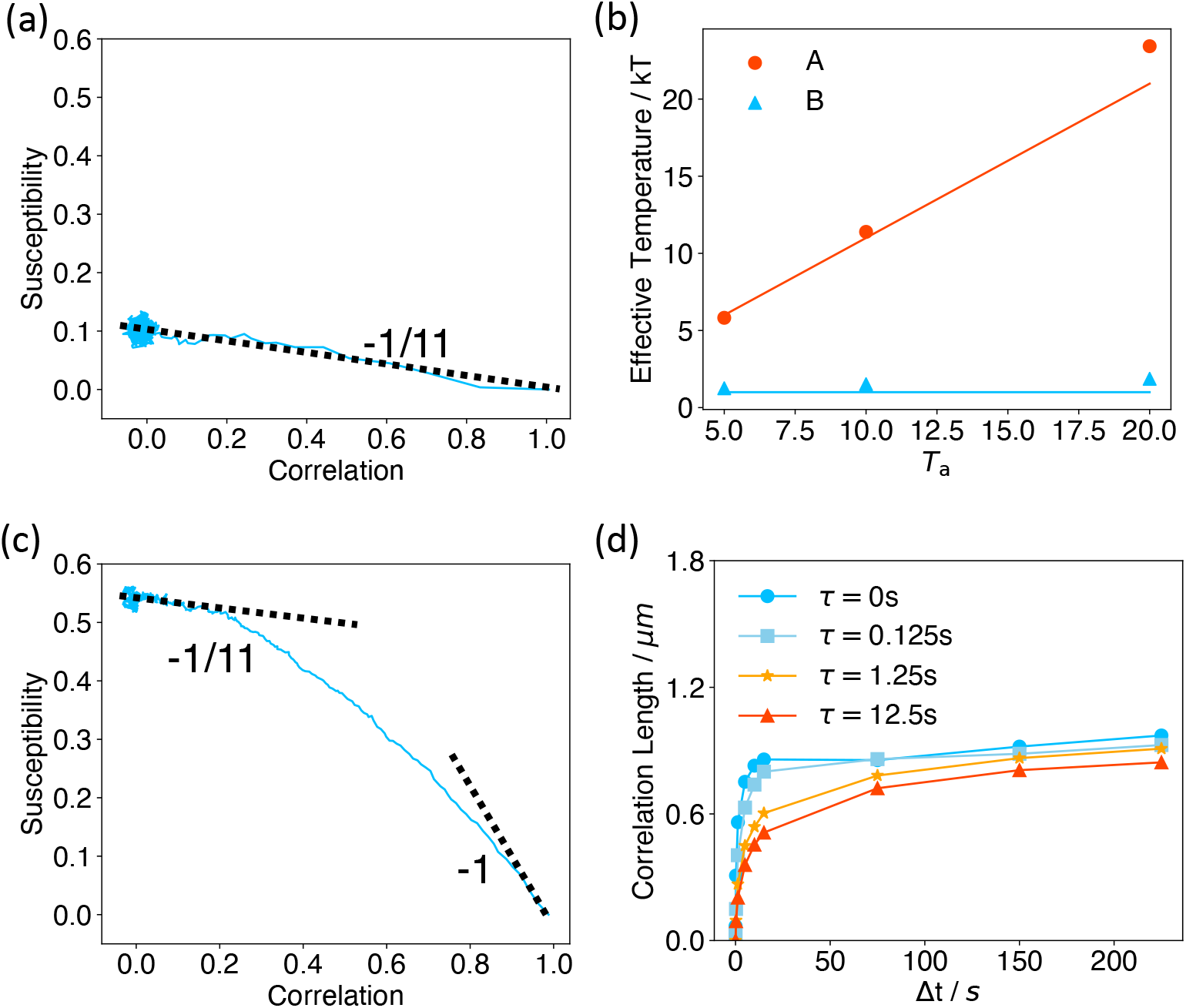
The effective temperature of active systems computed using the fluctuation-dissipation ratio. (a,c) Susceptibility as a function of correlation for the system with *T_a_* = 10*T*, *τ* = 0s (a) and *τ* = 1.25s (c). The dashed lines correspond to the thermal temperature – 1/*T* and the theoretical value – 1/(*T_a_* + *T*) for non-correlated active forces. (b) Effective temperatures of *A/B* compartments computed using the fluctuation-dissipation ratio for *T_a_* = 5,10, and 20T at *τ* = 0s are shown as dots. The red line corresponds to *T_a_* + *T*, while the blue line represents the constant *T*. (d) Correlation lengths as functions of time interval Δ*t* for different correlation time *τ* of the colored noise.

For correlated noises with nonzero *τ*, we found that the systems exhibit two timescales (Figure 5c and S11). At smaller timescales, the susceptibility-correlation plot for A compartments is approximately linear with a slope close to – 1/*T*, indicating that the system is in thermal equilibrium. At much longer timescales, the effect of non-equilibrium motors dominates, and the plot again becomes linear but with a different slope that approaches – 1/(*T_a_* + *T*). Furthermore, the transition regime between the two timescales increases with *τ* (Figure S11), suggesting that the system stays in the thermal temperature longer at larger *τ*. Therefore, for the same Δ*t*, larger *τ* leads to a lower effective temperature and the decreased correlation length seen in Figure 4d. The appearance of a higher effective temperature at longer timescales has indeed been predicted in prior simulation and theoretical studies. ^94,99,100^ As in the non-correlated noise, we anticipate that the higher effective temperature might lead to an increase in the correlation length at longer timescales. Indeed, we found that while *l_c_*(Δ*t*) has plateaued after around 15s for *τ* = 0, it continues to rise for larger *τ*. The value *l_c_*(Δ*t*) at different *τ* almost converges to the same value for Δ*t* = 225s (Figure 5d and S12).

## Conclusions and Discussion

We studied the impact of non-equilibrium activities on the structure and dynamics of the human genome. We found that activities such as transcription that are restricted to A compartments significantly influence the phase separation of euchromatin and heterochromatin. Heterochromatin is driven more towards the nuclear envelope, leaving more room in the interior for euchromatin. This effect can be largely understood from the presence of a temperature gradient. The impact on phase separation differs from those conferred by passive interactions, which stabilize the opposite trend by moving heterochromatin towards the nuclear interior, giving rise to the inverted nucleus.

Non-equilibrium phase separation between active and inactive particles has been shown to be sensitive to density. ^48,101^ We repeated the active force simulations in larger nuclei to examine the impact of density on genome organization. Figure S13 shows that the preferential localization of B compartments towards the nuclear envelope becomes less apparent in these simulations. We note that, for stem cells and cancer cells, which both have larger volumes, LADs, in fact, are less localized towards the periphery. ^102,103^ Though a disruption in lamin expression may cause the displacement of LADs from the nuclear envelope,^65,104–107^ the non-equilibrium mechanisms could contribute to the changes in LADs localization as well.

We note that the model presented here is still an approximation of the cell nucleus. Additional factors not accounted for in our model, including hydrodynamic effects, ^84^ vectorial forces, ^72^ and deformability of the nucleus envelope,^108^ could further enhance the spatial correlation length. Including these factors will be interesting in the future for building more quantitative models of the genome.

## Methods

### Computational model of the human genome

We followed the simulation strategy outlined in Ref. 17 to study the diploid human genome. The model explicitly represents the 46 chromosomes as beads-on-spring polymers confined in a sphere depicting the nucleus envelope. Based on the contact maps of Hi-C data from GM12878 cells, we designated each one of the coarse-grained one MB size beads as compartment *A, B*, or centromeric *C*. This assignment results in 2424 beads as compartment *A* and 3130 as *B*.

We simplified the energy function introduced by Qi et al. ^17^ to focus on the effect of non-equilibrium motors on genome organization. In particular, we enforced a common set of parameters for non-bonded interactions within the same and across different chromosomes. This setup provides a convenient system to explore molecular factors beyond simple passive interactions in positioning chromosomes inside the nucleus. Furthermore, we switched from a soft-core potential to the Lennard-Jones potential for non-specific interactions between genomic loci to avoid chain crossing. The soft-core potential was introduced originally to facilitate conformational sampling. However, the two potentials produce quantitatively similar trends for both dynamical and structural results (Figure S14). Detailed expressions of the energy function can be found in the Supporting Information.

We also adjusted the radius of the spherical confinement from 19.69 to 13.0 *σ*, where *σ* is the unit length and represents the diameter of a 1MB long genomic segment. The original confinement size was chosen to produce a DNA volume fraction of 0.1. Assuming a cell nucleus of 10 *μ*m in diameter, we estimated *σ* as 254 nm using the original confinement size. However, for genomic regions of one MB in length, the average size determined via superresolution imaging^109^ is larger than this value. Therefore, we rescaled the confinement size such that the estimated value for *σ* = 385 nm better matches experimental measurements.

### Details of dynamical simulations

We implemented the genome model in the LAMMPS software^110^ to carry out Brownian dynamics simulations. Without active forces, the equation of motion for particle i is defined as

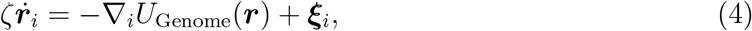

where *ζ* is the friction coefficient and *U*_Genome_(***r***) is the energy function of the genome defined in Eq. S1. The random force term ***ε**_i_* has zero means and satisfies the fluctuation-dissipation theorem as 〈***ε_i_***(*t*)***ε**_j_*(*t*′)〉 = 6*k_B_Tζδ_ij_δ*(*t* – *t*′).

Active forces were implemented by adding an additional noise term, ***f**_i_*, to the right hand side of Eq. 4. For *τ* = 0, the correlation function for ***f**_i_* as defined in Eq. 1 reduces to *6k_B_T_a_ζδ_ij_δ*(*t* – *t*′). Therefore, the two noise terms can be combined together to produce an effective equilibrium model with *T*_eff_ = *T* + *T_a_*. For non-zero *τ*, we treat the active force ***f**_i_* as dynamical variables that evolve with the following equation

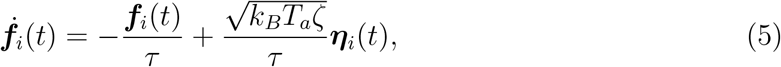

where ***η**_i_*(*t*) are Gaussian white noises that satisfy 〈(***η**_i_*(*t*) · ***η**_i_*(*t*′)〉 = 6*δ*(*t* – *t*′). Both Eq. 4 and 5 were integrated with the modified Euler-Maruyama algorithm. ^111^

Dynamical simulations were initialized with an equilibrium structure obtained from a long-timescale trajectory performed in our previous study.^17^ We carried out additional 5 × 10^6^ step-long equilibration simulations before switching to production runs that lasted for 2 × 10^7^ steps. The second half of production trajectories were divided into five non-overlapping blocks to provide independent estimates for variables of interest. The standard deviations of these estimates were shown as error bars of the mean values.

All simulations were performed with reduced units. We estimated the length unit *σ* = 385nm, the energy unit *∈* = *k_B_T*, where *T* is the room temperature. The time unit *τ_B_* = 0.125s was determined such that the diffusion constant of coarse grained particles match the experimentally determine values 10^−2^*μ*m^2^/s.^112–114^

## Acknowledgement

This work was supported by the National Institutes of Health grant R35GM133580.

## Supporting Information Available

- Details of the diploid Human genome model parameterized with Hi-C data, Brownian dynamics simulations, and numerical analysis, and supplementary figures.

## Notes

### Competing Interest Statement

The authors have declared no competing interest.

